# A functional medicine and food homology composition discovery based on disease-related target and data mining against cardiac remodeling

**DOI:** 10.1101/2024.01.14.575612

**Authors:** Dan Xiao, Runze Li, Xiaoqing Qin, Jinhai Feng, Denis Baranenko, Liudmila Natdochii, Yingyu Zhou, Jicheng Liu, Yan Lin

**Author notes:** Corresponding author to Yan Lin: Department of Scientific Research, School of Basic Medicine, Qiqihar Medical University, 333 bukui Street, Jianhua District, Qiqihar, Heilongjiang 161006, China.

## Abstract

**Background:** Medicine and food homological (MFH) products exhibit enhanced safety and tolerability, minimizing notable side effects, making them pivotal for prolonged use in cardiovascular diseases. This study aims to identify functional compounds in MFH based on cardiac remodeling-related target, employing reliable, comprehensive, and high-throughput methods.

**Methods:** By bioinformatics and *in vivo* verifications, we initially investigated the key target in the progression of cardiac remodeling. Subsequently, we performed molecular docking among medical homology compound database (MHCD), and then performed drug-likeness evaluations to recognize functional component based on disease-related target. Pharmacological verifications and data mining including cardiac and medullary transcriptomics, neurotransmitter metabolomics, resting-state functional magnetic resonance imaging (rs-fMRI), and correlationship analysis were utilized to define the benefical effects of MFH functional components, as well as its in-depth mechanims.

**Results:** The critical roles of oxidative stress and the key target of NRF2 in cardiac remodeling were discovered, and β-ecdysterone was screened as the most promising NRF2 enhancer in MHCD. Dose-dependent efficacy of β-ecdysterone in countering oxidative stress and ameliorating cardiac remodeling were then verfied by *in vivo* and *ex vivo* experiments. By data mining, the crosstalk mechanism between cardiac remodeling and neuromodulation was identified, and further unveiled *Slc41a3* as a potential key factor influenced by β-ecdysterone. Additionally, β-ecdysterone mitigated increases in norepinephrine (NE) and its metabolites DHPG in the sympathetic nerve center hypothalamic paraventricular (PVN), as indicated by rs-fMRI. Cardiac and medullary transcriptomes revealed central-peripheral regulation signaling pathways during cardiac remodeling with the involvement of core gene of *Dhx37*.

**Conclusions:** Our study identified β-ecdysterone as a natural MFH functional compound countering cardiac remodeling by targeting NRF2 elevation. It elucidates crosstalk between cardiac remodeling and neuromodulation, facilitating precise drug screening and mechanistic insights, providing substantial evidence for β-ecdysterone application and molecular mechanisms in cardiovascular diseases.

## Introduction

Natural product has more than 2,000 years of history and has gained widespread clinical applications. However, their explicit role in preventing and treating cardiovascular disease remains unclear due to a lack of sound scientific evidence^1^. Compared to synthetic drugs, natural active compounds often exhibit better safety and tolerability. This is attributed to their origin in nature, making them closer to the biological systems of the human body and causing relatively fewer noticeable side effects^2^. This advantage renders natural active compounds particularly important in long-term, particularly in the treatment of cardiovascular diseases^3^. Many natural bioactive compounds have been found to have protective effects on cardiovascular diseases. For example, flavonoids (such as soy isoflavones and anthocyanins) and polyphenols (such as catechins and anthocyanins) have positive effects on cardiovascular health. These compounds exhibit multiple actions, including antioxidant, anti-inflammatory, antiplatelet aggregation, and lipid-regulating effects, which can reduce the risk of cardiovascular diseases and improve cardiac function^4–6^. Cardiac remodeling is a key step in the development of heart failure and closely related to the prognosis of cardiovascular diseases^7^. Baicalin^8, 9^ and Danshen^10^ and their active compounds, such as resveratrol^11^ and emodin^12^, have been found to have anti-cardiac hypertrophy and anti-cardiac fibrosis effects, which can alleviate abnormal changes in cardiac structure and improve cardiac function. Notably, the newly defined sub-classification in natural product, medicine and food homology (MFH), offer notable benefits into a balanced diet not only aids in the prevention of neurodegeneration^13^, inflammation^14^, and aging^15^, etc., but also promotes overall well-being^16^. Based on the inherent properties of antioxidant and anti-inflammatory effects, our study first focused on their crucial role against cardiac remodeling, searching for the functional compounds of MFH to maintaining optimal cardiovascular system function.

The general limitations in the study of MFH primarily revolve around the absence of systematic natural product screening processes and drug similarity assessments^17^. While MFH holds potential therapeutic value, its intricate chemical structures and diversity necessitate large-scale screening and evaluation to discern its effects on cardiovascular diseases. Computational aided drug design (CADD) approaches offer an effective means of evaluating the absorption, distribution, metabolism, excretion, and toxicity characteristics of natural products and their active ingredients, enabling high-throughput and rapid screening^18, 19^. Thus, in our quest to identify a functional MFH composition with anti-cardiac remodeling properties suitable for long-term use, we employed a combination of bioinformatic tools, CADD, and pharmacological investigations.

Additionally, exploring the crosstalk between central and peripheral mechanisms in cardiovascular diseases is crucial for gaining a comprehensive understanding of overall regulation and interactions within the cardiovascular system. Hence, our current study aimed to delve into the underlying mechanisms, particularly the central-peripheral interactions, and established a scientific research methodology for the study of MFH-cardiac remodeling-crosstalk mechanisms.

## Methods

(Detailed methods were described in supplemental methods)

### Bioinformatics analysis

To identify cardiac remodeling-related genes, we employed two comprehensive and widely recognized databases: DisGeNet (https://www.disgenet.org/) and GeneCards (https://www.genecards.org/)^20^. String (https://string-db.org/) and Cytoscape (https://www.omicshare.com/tools) to define the key regulatory genes during cardiac remodeling and oxidative stress.

UCSC provided *Slc41a3* and *Dhx37* promoter sequences, and PROMO database predicted their transcription factors with 0% fault tolerance.

Open Knowledge Map with Pubmed database and keywords "Dhx37" facilitated literature search and classification. See Figure S9 for the classified literature.

### Preparation of medical and food homology compound database (MFCD) library

A MFCD library, comprising 110 known MFH natural products, established from publications by the National Health Commission of the People’s Republic of China. Utilizing TCMSP and literature, we expanded MFCD to include 3907 medical homologous components. LigPrep and Epik programs were employed for ligand preparation, generating low-energy 3D structures for subsequent molecular docking using Glide.

### Molecular Docking and Dynamics Simulation: KEAP1-NRF2 Binding Site

The KEAP1-NRF2 binding site structure (PDB identifier: 3ZGC) was obtained from RCSB. LeDock facilitated protein-ligand docking, and Maestro visualized the results. Molecular dynamics simulations, covering system setup, equilibration, and analysis, were conducted using gromacs and charmm 36 force field. Root-mean-square deviation (rmsd) measured structural stability over 100 ns.

### Absorption, distribution, metabolism, and excretion (ADME) prediction

ADME characteristics of lead compounds were calculated for drug discovery and were performed by ADMETlab 2.0 platform (https://admetmesh.scbdd.com/).

### Animal and in vivo experimental design

Ethical approval for animal experiments was obtained from the Ethical Committees of Qiqihar Medical University, adhering to the US National Institutes of Health guidelines. Kunming mice were acquired, housed in controlled conditions, and acclimatized for a week. The mice were randomly divided into Control, Cardiac Remodeling, and Treatment groups. After anesthesia, mice underwent cardiac echocardiography for cardiac functional measurement. Immunohistochemistry staining of heart tissues were processed for hematoxylin-eosin (HE), Masson trichrome, and Sirius red staining.

### Cell culture and ex vivo experimental design

Cardiac fibroblasts and cardiomyocytes were isolated from neonatal mice and transfected with Nrf2 siRNA or control siRNA. Transfected cells were treated with AngII and β-ecdysterone, and subsequent cell number, viability, oxidative stress, inflammation, protein/RNA extraction was performed.

### Resting-state Functional Magnetic Resonance Imaging (rs-fMRI)

A separate cohort underwent T2-weighted MRI and rs-fMRI analysis. Image preprocessing was performed using MATLAB toolboxes. Dynamic patterns and temporal variability of intrinsic brain activity were characterized using DynamicBC toolbox.

### Cross-talk mechanism mining

Transcriptomics of cardiac and medullary tissues were performed by utilizing the Illumina sequencing platform. WGCNA analysis were then used to calculated the potential relationship of cardiac remodeling versus the transcriptomic data of cardiac and medullary tissues.

Cardiac neurotransmitter metabolomics were then conducted to analyze percentage composition of each metabolite within samples. Top 20 metabolites by abundance were displayed.

### Statistical analysis

All values were presented as mean ± S. E. M. Statistical comparisons were performed by Student’s t-test between two groups or one-way ANOVA for multiple comparisons, and p < 0.05 was considered to indicate a significant difference. Data were analyzed using the GraphPad Prism 7.0 software.

## Results

### β-ecdysterone potentially releases NRF2 via completely binding to KEAP1

Cardiac remodeling constitutes a pivotal pathological alteration in cardiovascular diseases, primarily distinguished by myocardial hypertrophy and myocardial fibrosis. To unravel the pivotal pathways implicated in cardiac remodeling, we leveraged the Gene Cards database to investigate "cardiac hypertrophy" and "cardiac fibrosis," conducting functional enrichment analysis on the shared genes (Fig. S1A & B). Our analysis pinpointed the involvement of pathways related to oxidative stress (Fig. S1B). Subsequently, utilizing the DisGenet database, we identified core genes associated with "oxidative stress" and recognized NRF2 as a central player in the oxidative stress pathway (Fig. S1C). To authenticate the alterations in oxidative stress and NRF2 during cardiac remodeling, we employed three established animal models: aortic constriction (TAC), isoproterenol administration (ISO), and L-NAME-induced hypertension. Through quantification of the oxidative stress marker MDA and NRF2 protein levels, we discerned the model most faithfully representing cardiac remodeling. The findings revealed a significant surge in oxidative stress levels and a concurrent downregulation of NRF2, a transcription factor impeding oxidative stress, in cardiac tissue undergoing cardiac remodeling. Notably, the most pronounced downregulation of NRF2 occurred in the ISO animal model (Fig. S1D and E). Consequently, we selected the ISO animal model for subsequent research and analysis.

NRF2 binds to its inhibitory factor KEAP1, leading to the recognition of ubiquitination sites and the degradation of the NRF2 protein. Therefore, our research is focused on this process to identify MFH composition that competitively bind to KEAP1, thereby releasing more NRF2 to exert antioxidant and stress response effects. Utilizing computer-aided drug design methods, we targeted the binding site of KEAP1-NRF2 (Fig. 1A, B) and screened candidate compounds from MFCD. The screening criteria were as follows: 1) The ADME prediction result scored 0 (STARS: 0), indicating that the compound meets all the criteria for drug-like properties. This compound exhibits excellent properties related to drug absorption, distribution, metabolism, and includes other characteristics such as hydrogen bond acceptors within the specified range. 2) Molecular docking score < -7.038^20^. 3) Using principles of drug-likeness such as Lipinski’s rule of five and Veber’s rule of three, compounds were screened with the criterion of no more than one violation. Based on this, 66 natural product molecules were selected (Table S1) for further assessment of drug-likeness. In the assessment of drug-likeness recorded in Traditional Chinese Medicine Systems Pharmacology (TCMSP) platform, we initially screened Molecular weight (MW) < 500, LogP < 5, Drug-likeness (DL) > 0.1 and Oral bioavailability (OB) > 20% (Table S2). Consequently, β-ecdysterone was found be the best MFH composition meeting all the screening criteria among 10 compositions. The properties of β-ecdysterone (Table 1) and its stability when binding to KEAP1 in molecular dynamics simulation within 100 ns (Fig. 1C, D) were performed using Schrödinger’s drug-likeness evaluation and molecular dynamics simulation. The results showed that β-ecdysterone has perspective drug-like potential with stable binding to KEAP1.

**Figure 1.**
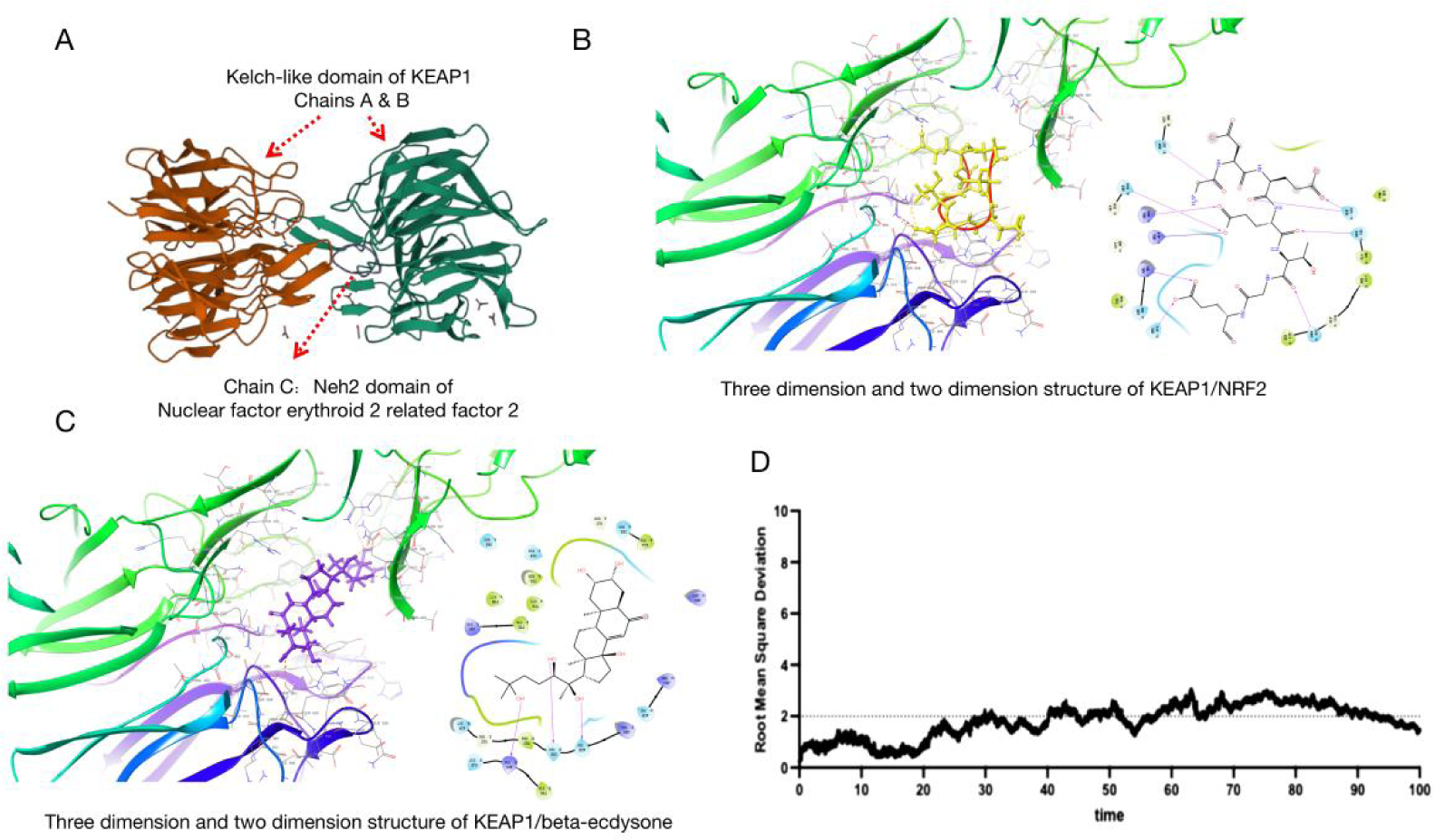
Using computer-aided drug design, competitive binding natural product β-ecdysterone was found for the binding site of NRF2 and KEAP1. **(A)** 3D protein structure of the binding site of NRF2 and KEAP1; **(B)** 3D and 2D representation of the binding of NRF2 to KEAP1 through molecular docking; **(C)** 3D and 2D representation of the binding of β-ecdysterone to KEAP1 at the binding site of NRF2 and KEAP1; **(D)** Evaluation of the binding between β-ecdysterone and KEAP1 using molecular dynamics simulation for a duration of 100 ns.

**Table 1.** Drug-likeness properties of β-ecdysterone.

| Permeability |  |  |
| --- | --- | --- |
| Log P | 1.75 | Recommendation: 1~3 |
| Log S | -2.78 | Recommendation: -1~-5 |
| Water Solubility (mg/mL) | 0.415 | Recommendation: 0.1~100 |
| Hydrogen Acceptor Count | 7 | Recommendation: $\leq 10$ |
| Hydrogen Donor Count | 6 | Recommendation: $\leq 5$ |
| Polar Surface Area ($\text{\AA}^2$) | 134.899 | Recommendation: 20~140 |
| Rotatable Bond Count | 5 | Recommendation: $\leq 10$ |
| Number of Rings | 4 | Recommendation: 3~6 |
| Molecular weight | 480.71 | Recommendation: 250~500 |
| Molar Refractivity | 129.74 | Recommendation: 40~130 |
| Fractional negative accessible surface area (FASA-) | 0.26 | Recommendation : $\leq 0.339$<br>(PMID: 23181938) |
| **Oral bioavailability** |  |  |
| Oral availability (%) | 44.23 | Recommendation: $\geq 20$ |
| Human Oral Absorption (%) | 55.180 | Class 2 |
| PCaco-2 | 56.299 | High gastrointestinal absorption |
| BBB permeant | -2.05 | Non-penetrating potential |
| **Metabolism and plasma stability** |  |  |
| $t_{1/2}$ (h) | 4.51 | |
| IP(eV) | 10.298 | Recommendation: 7.9~10.5 |
| EA(eV) | 0.417 | Recommendation: -0.9~1.7 |
| **Drug likeness evaluation** |  |  |
| Drug likeness (DL) from TCMSP | 0.82 | Recommendation: $\geq 0.1$ |
| Druglikeness of Lipinski | Yes | 1 violation: Hydrogen Donor Count equals 6 ($>5$) |
| Druglikeness of Veber | Yes |  |
| Druglikeness of Ghose | No | 2 violations: molecular weight equals 480.71 ($>480$), total atoms equals 78 ($>70$) |

### Pharmacological investigation of β-ecdysterone on cardiac remodeling

To gain insights into the effects of β-ecdysterone on cardiac remodeling, we initially administered ISO through subcutaneous injections to establish animal models, with or without β-ecdysterone treatment for a continuous period of 14 days. Subsequently, we observed significant elevations in cardiac functional parameters, including heart weight/tibia length (HW/TL), fractional shortening (FS), and ejection fraction (EF). In line with previous findings^21^, these changes in cardiac function and morphology are primarily attributed to compensatory remodeling stimulated by ISO overload. However, β-ecdysterone treatment mitigated the alterations induced by ISO on cardiac function and morphological dimensions in a dose-dependent manner (Fig. 2A-D). To specify, the echocardiography parameters revealed ISO-induced changes in cardiac left ventricle mass and cardiac function (Fig. 2A), heart volume, and heart weight index (Fig. 2B), as well as morphological and biomarker changes associated with cardiac remodeling (Fig. 2C-F). Notably, all these effects were mitigated by β-ecdysterone treatment, as illustrated in Figure 2.

**Figure 2.**
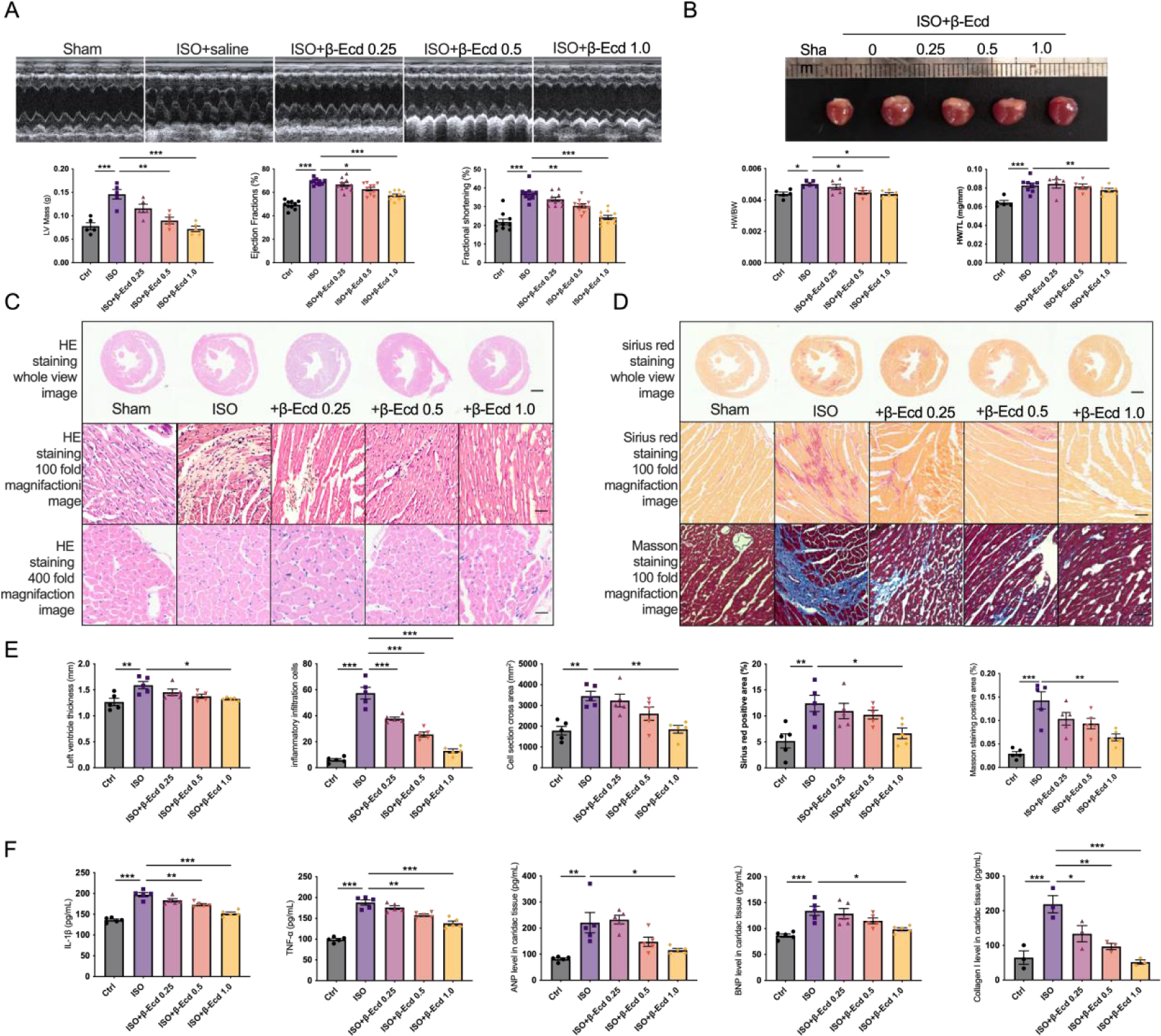
β-ecdysterone ameliorates the elevation of compensative cardiac function in ISO-treated mice. **(A)** According to the echocardiogram images and statistical results, the left ventricular weight (LV Mass), ejection fraction (EF) and fractional shortening (FS) was recorded, n = 5-10 in each batches, *p < 0.05, **p < 0.01, ***p < 0.001; **(B)** Whole heart images from mice are showed at the top with statistical results of the ratio between heart weight/body weight (HW/BW) and heart weight/tibia length (HW/TL). Mice subjected to ISO injection showed an increase in volume compared with controls and ameliorated by β-ecdysterone treatment in dose-dependent manner, n = 5-8 in each batch, *p < 0.05, **p < 0.01, ***p < 0.001; **(C)** Images of HE staining was performed to analysis of the morphology, heart wall thickness, inflammation, cardiomyocyte cross section area of heart tissue. From the top to bottom are presented as the whole view of heart tissue, which indicates the morphology and thickness of the heart wall (Scale bar equals 1 mm), 100 magnification, which indicates the inflammatory infiltration (Scale bar equals 50 µm) and 400 magnification, which indicates cardiomyocytes cross section area (Scale bar equals 12.5 µm); **(D)** Images of Sirius red staining and Masson staining to analysis the collagen composition of heart tissue. From the top to bottom are presented as the whole view of Sirius red staining (Scale bar equals 1 mm), 100 magnifications of Sirius red staining (Scale bar equals 50 µm) and 100 magnifications of Masson staining (Scale bar equals 50 µm). **(E)** Statistical data of (C) and (D) are presented, including (from left to right) heart wall thickness, inflammatory infiltration cells, cell section cross area, Sirius red staining positive area, and Masson staining positive area. n = 5 in each batch, *p < 0.05, **p < 0.01, ***p < 0.001; (F) Cardiac remodeling markers were detected by using ELISA assays, the parameters include (from left to right) inflammatory factors (IL-1β, TNF-α), cardiac hypertrophic markers (ANP, BNP), and cardiac fibrotic markers (Collagen I). n = 3-5 in each batch, *p < 0.05, **p < 0.01, ***p < 0.001.

Inflammation, driven by factors like IL-1β and TNF-α, is pivotal in diverse pathological processes. These widely studied inflammatory factors intensify diseases, forming a harmful cycle without intervention. Our study explores the potential anti-inflammatory role of β-ecdysterone in mice injected with ISO. Inflammatory infiltration was assessed through HE staining, as depicted in Figure 2C, revealing that β-ecdysterone indeed decreased ISO-induced inflammation in the left ventricle heart tissue in a dose-dependent manner. Furthermore, ELISA assays were employed to measure the inflammation markers IL-1β and TNF-α (Fig. 2F), yielding consistent results that affirm the anti-inflammatory effects of β-ecdysterone against ISO.

Combining these results, β-ecdysterone not only influences inflammatory infiltration but also modulates inflammatory factors, thereby ameliorating cardiac impairment caused by ISO overload. Based on the findings, we selected a dosage of 1.0 mg/kg β-ecdysterone for subsequent animal experiments.

### β-ecdysterone: A superior MFH compound for cardiac hypertrophy in comparison to fibrosis in ISO-induced cardiac remodeling

Cardiac remodeling is generally considered to involve two main processes: cardiac hypertrophy and cardiac fibrosis. Cardiac hypertrophy entails the enlargement and thickening of heart muscle cells, while myocardial fibrosis refers to the activation of cardiac fibroblasts into myofibroblasts, leading to excessive proliferation and the accumulation of collagen. In this study, we aimed to investigate the effects of β-ecdysterone on cardiac hypertrophy and cardiac fibrosis in ISO-induced cardiac remodeling using an *in vitro* experimental design.

Initially, we isolated cardiomyocytes and cardiac fibroblasts from neonatal mouse hearts through primary isolation and employed Angiotensin II (AngII, 10^-7^ M) *in vitro*. Subsequently, we assessed whether β-ecdysterone treatment affected cardiomyocytes, cardiac fibroblasts, or both. We initially explored the suitable, as well as the toxicity of β-ecdysterone concentrations for cardiomyocytes. Doses exceeding 80 μM β-ecdysterone resulted in a sharp decrease in the cell number and viability of cardiomyocytes. Consequently, we included concentrations no more than 80 μM and opted for 10-40 μM as an effective concentration in subsequent *in vitro* experiments (Fig. S2).

Concerning the anti-hypertrophic effects, β-ecdysterone significantly reduced the cross-sectional area (CSA) of cardiomyocytes in a dose-dependent manner, which had been enlarged by AngII treatment; The hypertrophic marker, BNP expression level, was detected by immunofluorescent staining and was significantly downregulated in the β-ecdysterone-treated group in a dose-dependent manner compared to the AngII-treated group; Regarding NRF2, the basis of our MFCD selection, the expression pattern was rescued against AngII stimulation by β-ecdysterone treatment in a dose-dependent manner; Concerning the antioxidant stress response mediated by NRF2, we assessed various parameters, including reactive oxygen species (ROS), malondialdehyde (MDA), superoxide dismutase (SOD), catalase (CAT), and glutathione peroxidase (GSH-Px) levels. The results showed that AngII remarkably elevated oxidative stress levels, whereas β-ecdysterone treatment ameliorated this effect in a dose-dependent manner (Fig. 3 A & B).

**Figure 3.**
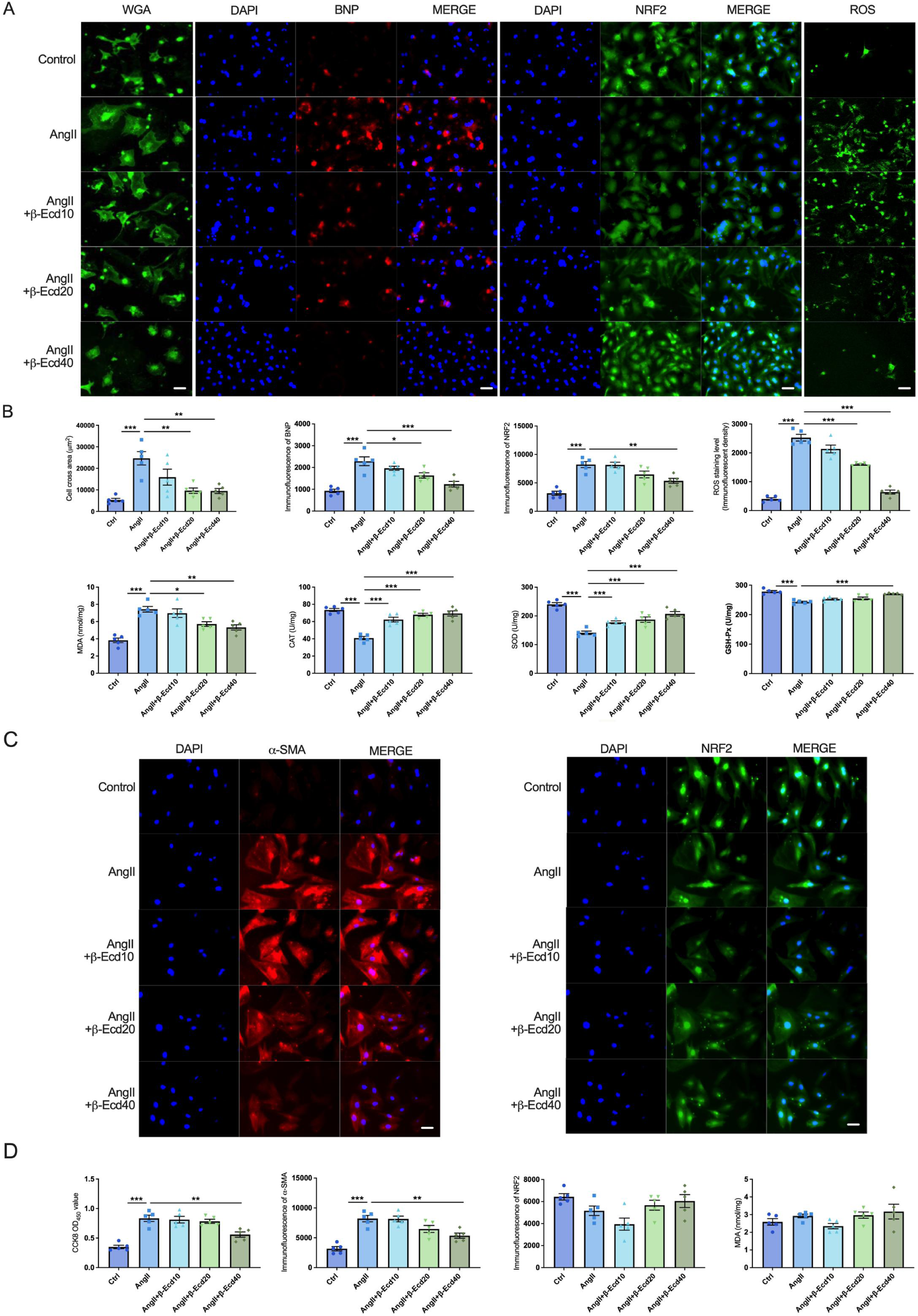
β-ecdysterone dose-dependently abolished AngII induced hypertrophy in cardiomyocytes. **(A)** Images of immunofluorescent staining in cardiomyocytes. From left to right are the images of WGA, DAPI & BNP, DAPI & NRF2, and ROS detection. Scale bar equals 12.5 µm; **(B)** In cardiomyocytes analysis, statistical data from (A) at the top line and oxidative stress analysis at the bottom line. At the top line of panels (from left to right, in sequence) are cell section area (CSA), immunofluorescent intensity of BNP (per cell), immunofluorescent intensity of NRF2 (per cell), and immunofluorescent intensity of ROS staining. At the bottom line of panels (from left to right, in sequence) are malondialdehyde (MDA), catalase (CAT), superoxide dismutase (SOD), and glutathione peroxidase (GSH-Px) levels. n=5 in each batch, *p < 0.05, **p < 0.01, ***p < 0.001; **(C)** Images of immunofluorescent staining in cardiac fibroblasts. From left to right are the images of DAPI & BNP and DAPI & NRF2. Scale bar equals 12.5 µm; **(D)** In cardiomyocytes analysis, statistical data of CCK-8 analysis, immunofluorescent intensity of α-SMA (per cell), immunofluorescent intensity of NRF2 (per cell) and oxidative stress analysis of MDA. n=5 in each batch, **p < 0.01, ***p < 0.001.

Concerning the anti-cardiac fibrosis effects (Fig. 3C & D), the 40 μM β-ecdysterone dose demonstrated inhibitory effects on cardiac fibroblast viability and their transformation into myofibroblasts, as evidenced by the reduced expression of α-SMA; Notably, while β-ecdysterone effectively attenuated the activation of cardiac fibroblasts, the expression of NRF2 was not adversely affected by AngII stimulation, and there was no observable increase in oxidative stress.

Regarding the anti-cardiac fibrosis effects (Fig. 3C & D), 1) the 40 μM β-ecdysterone dose indeed exhibited inhibitory effects on cardiac fibroblast viability and the transformation into myofibroblasts, as evidenced by the detection of α-SMA expression; 2) However, although β-ecdysterone effectively attenuated the activation of cardiac fibroblasts, the expression of NRF2 was not impaired by AngII stimulation, nor was there an increase in oxidative stress.

Considering these findings collectively, we boldly hypothesize that β-ecdysterone could alleviate cardiac hypertrophy through the regulation of NRF2-mediated oxidative stress, aligning with our initial plan for selecting natural MFH compounds. However, it is noteworthy that high doses of β-ecdysterone may impact cardiac fibroblast activation through specific mechanisms.

Mitochondrial dysfunction, attributed to cellular oxidative stress response as the source of reactive oxygen species, was identified as a key factor. Subsequently, we delved into the impact of β-ecdysterone on mitochondria. We assessed the functionality of respiration complexes I-IV and employed the JC-1 assay. As anticipated, AngII disrupted mitochondrial functions in cardiomyocytes, while treatment with β-ecdysterone ameliorated this disturbance (Fig. S3).

### NRF-2 serves as a mediator for the beneficial effects of β-ecdysterone on hypertrophic cardiomyocytes

As a central transcription factor in the cellular antioxidant stress response, NRF2 activates the endogenous antioxidant pathway, playing a pivotal role in ROS-induced antioxidant protein expression. To confirm whether NRF2 mediates the beneficial effects of β-ecdysterone against AngII in cardiomyocytes, we employed siRNA NRF2 to knock down NRF2 expression in primary isolated cardiomyocytes. The effectiveness of the transfection was verified (Fig. S4). Subsequently, we examined the previously mentioned beneficial effects of β-ecdysterone, encompassing anti-hypertrophy, anti-oxidative stress, and anti-mitochondrial dysfunction *in vitro*.

The downregulated expression of NRF2 due to siRNA transfection hindered the beneficial effects of β-ecdysterone, as evidenced by the compromised anti-hypertrophic effects (Fig. 4A-F), diminished anti-oxidative stress response (Fig. 4G-K), and impaired mitigation of mitochondrial dysfunction (Fig. 4L). These results strongly suggest that NRF2 is a crucial mediator of the functional effects of β-ecdysterone on cardiomyocytes.

**Figure 4.**
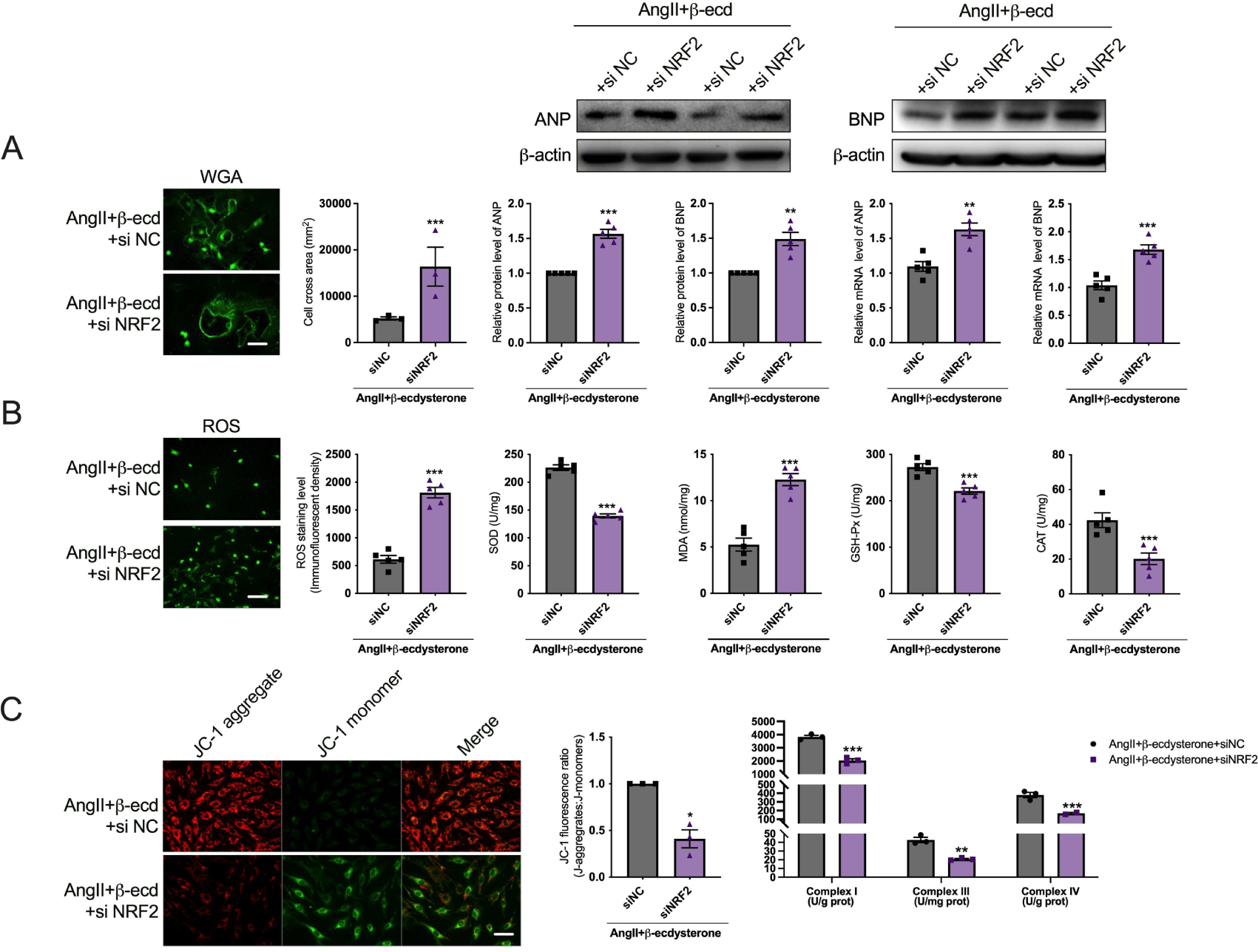
NRF2 mediates the effects of β-ecdysterone on cardiac dysfunction *in vitro*. **(A)** Verification the key mediated-function of NRF2 on cardiac hypertrophy in primary isolated cardiomyocytes during β-ecdysterone treatment. siNC represents primary isolated cardiomyocytes transfected with siRNA negative controls, siNRF2 represents this transfection with siRNA NRF2. From left to right, in sequence, present as the images with its statistical results of WGA staining, western blotting images with its statistical results of ANP & BNP, and mRNA levels of ANP & BNP. n=3-5, \*\**p* < 0.01, \*\*\**p* < 0.001; **(B)** The effect of siNRF2 transfection on oxidative stress were defined by ROS immunofluorescent staining, SOD, MDA, GSH-Px, and CAT assays. n=5, \*\*\**p* < 0.001; (C) Mitochondrial complex I, III & IV and JC-1 assay were determined by their kits to examine the effect of siNRF2 transfection on mitochondrial functions, n=3, \**p* < 0.05, \*\**p* < 0.01, \*\*\**p* < 0.001.

### 2.5 Transcriptome analysis of cardiac tissue: Unveiling the impact of β**-**ecdysterone on cardiac remodeling processes

Regarding β-ecdysterone, a natural MFH product identified in our MFCD screening with anti-hypertrophic potential, transcriptome analysis offers in-depth insights into its mechanisms of action and has substantial implications for guiding drug development and personalized therapy. Through differential gene expression and functional pathway analyses, we explored cardiac remodeling and the influence of β-ecdysterone administration on the cardiac transcriptome. Differential expression genes (DEGs) influenced by ISO treatment were found to be abundant in extracellular matrix organization and metabolic functions, such as propanoate metabolism and carbon metabolism. This observation aligns with the physiological function of ISO in activating cardiac β-receptors (Fig. 5A & B). Notably, following β-ecdysterone administration, several pathways associated with neurotransmitters were altered, including chemical synaptic transmission, the neuropeptide signaling pathway, and neuroactive ligand-receptor interaction (Fig. 5C & D). Interestingly, considering that β-ecdysterone was predicted to be unable to cross the blood-brain barrier (Table 1, BBB = -2.05), it strongly suggests an indirect impact of β-ecdysterone on the central nervous system.

**Figure 5.**
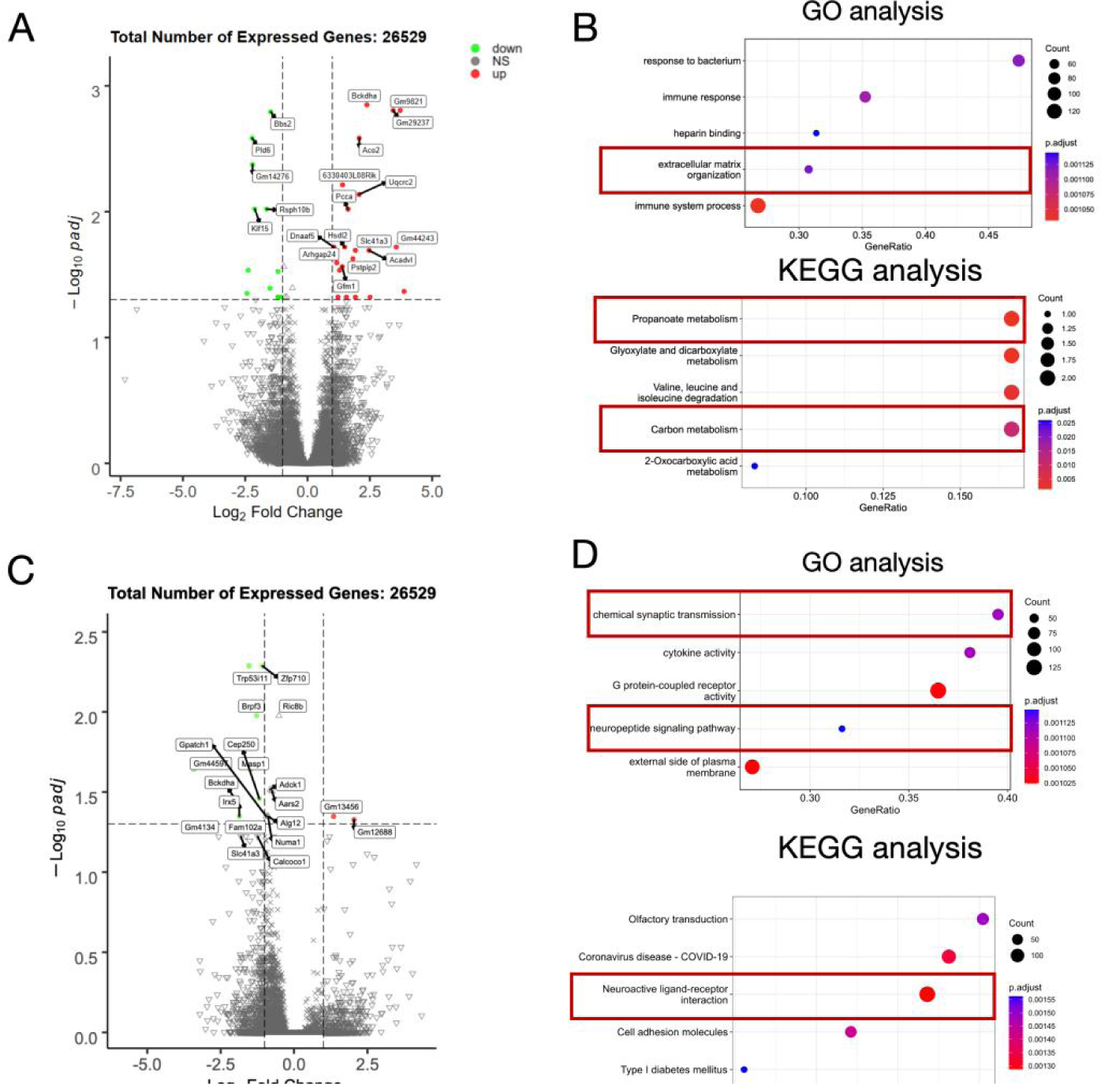
Transcriptomics Data of differential expression genes, function signaling pathways enrichment analysis in left ventricle of heart tissue. **(A)** In comparison of ISO with control group, differential expression genes (DEGs) were analyzed and calculated by Volcano plot. DEGs were then filtered by -log10P adj > 1.3, log2 Fold Change > ± 1. Totally genes were 21 genes upregulated whereas 12 genes were downregulated; **(B)** Functional enrichment of GO analysis (Top 5) and signaling pathway enrichment of KEGG analysis (Top 5) were presented to demonstration the potential mechanism of DEGs in comparison of ISO with control group. The red box represents functional/signal pathways associated with ventricular remodeling; **(C)** In comparison of ISO+β-Ecd with ISO group, differential expression genes (DEGs) were analyzed and calculated by Volcano plot. DEGs were then filtered by -log10P adj > 1.3, log2 Fold Change > ± 1. Totally genes were 8 genes upregulated whereas 2 genes were downregulated; **(D)** Functional enrichment of GO analysis (Top 5) and signaling pathway enrichment of KEGG analysis (Top 5) were presented to demonstration the potential mechanism of DEGs in comparison of ISO+β-Ecd with ISO group. The red box represents functional/signal pathways associated with neurotransmitter-related signaling pathways.

Furthermore, for a statistical analysis of the overlap between genes exhibiting significant changes following ISO stimulation and those altered after β-ecdysterone administration, we employed a Venn diagram to calculate the intersection of two distinct sets of differentially expressed genes (Fig. S5A & Table S3), excluding pseudo-genes (*Rps18-ps4*, *Zfp708*, *Tmem171*, *Gm43544*, *Gm15710*, *Gm1291*8, *Gm6058*), non-human heart specific genes (*Trpc2*), our attention was directed to the gene we finally focused on the gene of *Slc41a3*, primarily involved in ion transport.

In our transcriptome analysis, we observed significant upregulation of *Slc41a3* in ISO-treated heart tissue, and conversely, downregulation when treated with β-ecdysterone. To validate these transcriptomic findings, we utilized real-time PCR and immunohistochemistry to localize and quantify the expression of *Slc41a3* in cardiac tissue. Consistent with the transcriptomic results, *Slc41a3* mRNA and SLC41A3 protein levels exhibited high expression in the ISO group, while β-ecdysterone (at a dose of 1.0 kg/mg) suppressed their expression (Fig. S5B-D).

### β-ecdysterone modulates nervous systems in cardiac remodeling

Based on the preceding results, we have discerned the intervention effect of β-ecdysterone on the alterations in pathways related to neurotransmitters in the left ventricle of heart tissue. Considering the regulation of cardiac tissue by the sympathetic and parasympathetic nervous systems to maintain functional homeostasis (Figure S6A), we initially hypothesized that β-ecdysterone might recover this balance against ISO treatment. We initially hypothesized that β-ecdysterone might enhance central nervous system feedback by upregulating the activity of the parasympathetic nervous system. We measured the levels of the sympathetic neurotransmitter norepinephrine (NE) and the parasympathetic neurotransmitter acetylcholine (Ach) in the left ventricle of heart tissue. However, the results obtained were different from our initial expectations. The levels of the sympathetic neurotransmitter NE significantly decreased following β-ecdysterone intervention, whereas the changes in acetylcholine levels were not pronounced (Fig. S6B, C).

To elucidate the specific reasons behind this and further validate the result, we conducted neurotransmitter metabolomics profiling for an additional 33 common neurotransmitters. The results, presented in Figure 6, utilized a series of statistical methods to identify neurotransmitters exhibiting significant differences among various groups (Fig. 6A-D). We then pinpointed five neurotransmitters associated with cardiac remodeling parameters (Fig. 6E & F), including heart weight (HW/BW), cardiac functional index (EF, FS), left ventricle hypertrophic index (LVHI), cardiac hypertrophic biomarkers (ANP, BNP), inflammatory factors (IL-1β), and oxidative stress (MDA), with a prominent role observed for 3,4-dihydroxyphenylethylene glycol (DHPG) (Fig. 6G). DHPG, a metabolic product derived from dopamine, showed a high correlation with norepinephrine metabolic levels (Fig. S7) and has been demonstrated to activate sympathetic activity. Importantly, β-ecdysterone significantly reduced the levels of DHPG in cardiac tissue, confirming its inhibitory effects on sympathetic activity.

**Figure 6.**
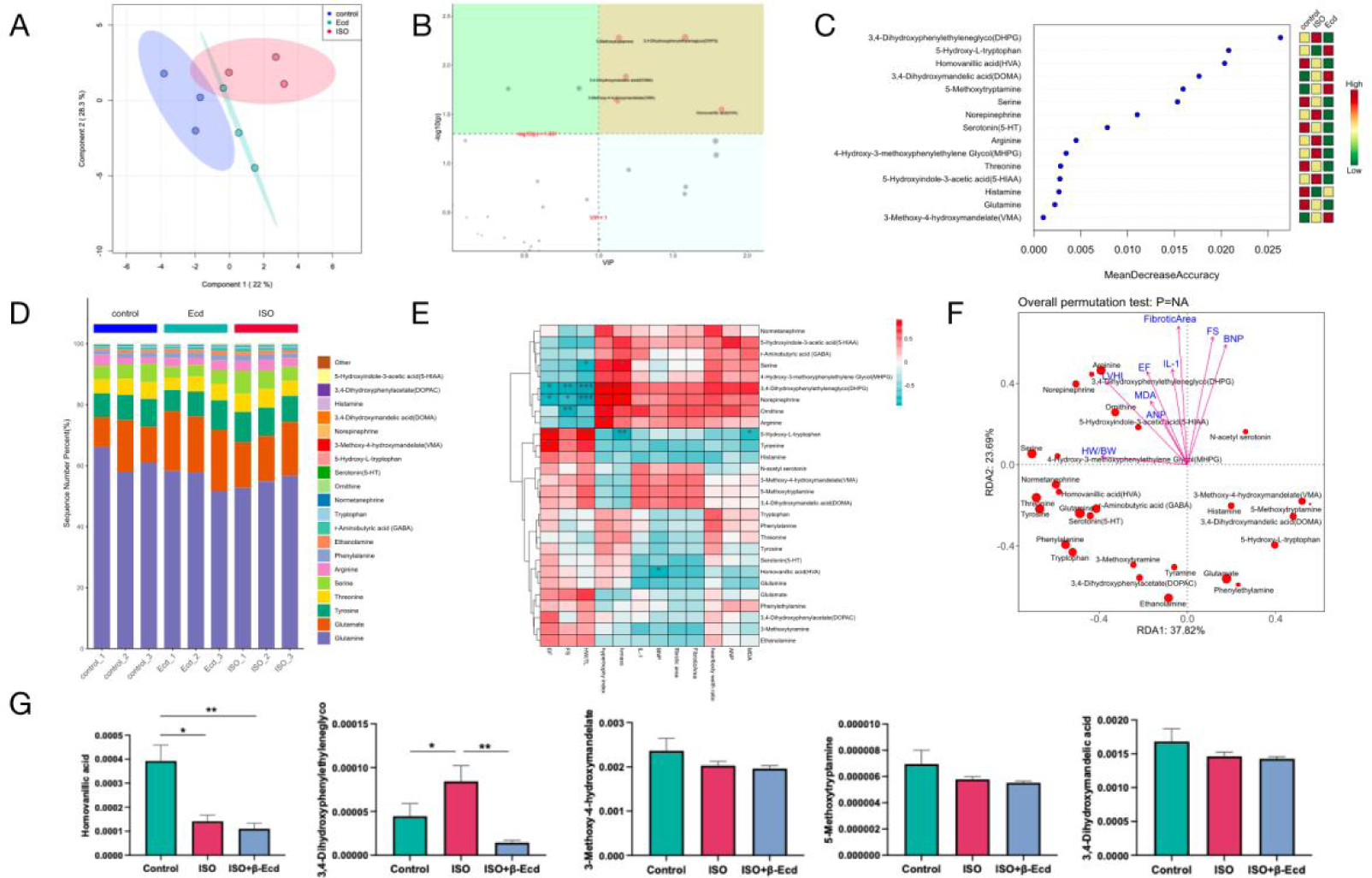
Neurotransmitter metabolomics profiling for other 33 common neurotransmitters in left ventricle of heart tissue. **(A)** Principal Component Analysis of (PCA) samples, n=3 in each group. **(B)** Value Importance in Projection (VIP) was performed. The metabolites highlighted in the yellow region are those with a corrected p-value less than 0.05 and a VIP score greater than 1. to identify the metabolites with a VIP score greater than 1 contribute significantly to discriminant analysis, which exhibit substantial differences between groups. **(C)** Random forest analysis was used with value of "mean decay accuracy" and "mean decay Gini" to measure the importance of a metabolite in distinguishing groups in random forest. The greater the two values, the greater the importance of metabolites in random forests; **(D)** Percentage content bar chart is used to calculate the top 20 content of metabolites, determining the percentage content of each metabolite within each sample. The results are then visualized using a stacked bar chart, allowing for a straightforward comparison of differences in metabolite composition structure between groups; **(E)** Correlation analysis between the characterization of left ventricular remodeling (Cardiac functional index: EF, FS; Cardiac remodeling index: HW/BW & left ventricle hypertrophic index (LVHI); Cardiac hypertrophic index: ANP & BNP; Cardiac fibrosis index: value of fibrotic area; Cardiac inflammation: level of IL-1β; Oxidative stress level: level of MDA) and neurotransmitters were performed and then presented as heat map. *p < 0.05, **p < 0.01, ***p < 0.001; **(F)** By employing regression relationship between the characterization of left ventricular remodeling (blue font) and neurotransmitters (red plot) to retain the explainable variations. Dimensionality reduction is applied, and their correlation is indicated using angle measurements; **(G)** Neurotransmitters of Homovanillic acid (HVA), 3,4- Dihydroxyphenylethyleneglyco (DHPG), 3-Methoxy-4-hydroxymandelate (VMA), 3-Methoxytyramine, 3,4-Dihydroxyphenylacetate (DOPAC) were presented as bar charts. Statistical analysis is performed through inter-group comparisons using the one-way ANOVA method. n=3 in each group, *p < 0.05, **p < 0.01.

In summary, we are delighted by the discovery that β-ecdysterone not only targets the key oxidative stress regulator NRF2, exerting an inhibitory effect on oxidative stress and enhancing cardiac remodeling, but also inhibits sympathetic nervous system activity. This dual action effectively alleviates the damage caused by excessive sympathetic activation induced by ISO.

### Crosstalk mechanisms of cardiac central-peripheral interaction in β-ecdysterone-ameliorated Cardiac remodeling

To broaden the understanding of the functional scope of β-ecdysterone, we employed resting-state functional magnetic resonance imaging (rs-fMRI). Through a comprehensive analysis with particular emphasis on the activity of the paraventricular nucleus (PVN), a pivotal component of the sympathetic nervous system, intriguing results are depicted in Figure S8. In ISO groups administered with and without β-ecdysterone, the activity level was assessed by calculating the amplitude of low-frequency fluctuations (ALFF) values. Through comparison with a brain atlas, we identified the PVN within the red circles and discovered that administration of β-ecdysterone significantly suppressed the ALFF values compared to the ISO group. Thus, we obtained direct evidence supporting the inhibitory role of β-ecdysterone in the central sympathetic nervous system (PVN).

Additionally, the parasympathetic fibers of the vagus nerve originate from the dorsal nucleus of the vagus nerve within the medulla oblongata, while the preganglionic fibers of the sympathetic nervous system originate from the central sympathetic nucleus within the medulla oblongata. Given the crucial role of medulla oblongata as the cardiovascular center, this section focuses primarily on whether β-ecdysterone further affects the medulla oblongata, responsible for maintaining the balance between the sympathetic and parasympathetic nervous systems.

Due to the limited capabilities of our rs-fMRI to detect medullary activity, we opted to extract medullary tissue from mice and applied transcriptome analysis to explore the effects of β-ecdysterone intervention from a genetic perspective. The results, as shown in Figure 7, revealed that ISO influenced the energy metabolism, while β-ecdysterone affected the extracellular matrix collagen remodeling in the medullary tissue, showing a positive correlation with cardiac relaxation. These findings indicate the involvement of β-ecdysterone in the medulla oblongata regulation of cardiac structure and function, consistent with our Figure 1 & 2, implying the influence of β-ecdysterone on the neuro-regulation and feedback mechanism from a central-to-peripheral perspective.

**Figure 7.**
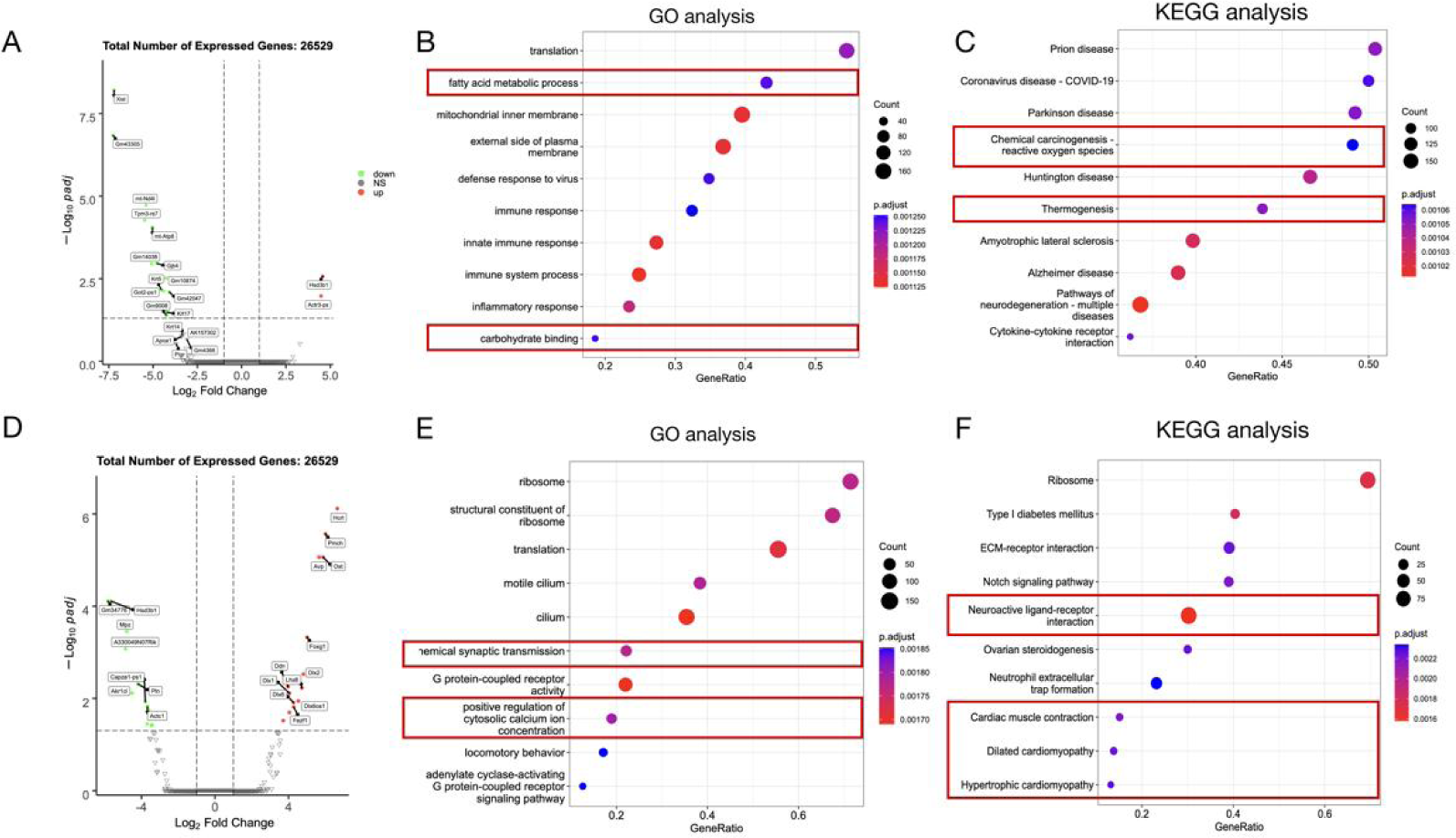
Transcriptomics Data of differential expression genes, function signaling pathways enrichment analysis in medullary tissue. **(A)** In comparison of ISO with control group, differential expression genes (DEGs) were analyzed and calculated by Volcano plot. DEGs were then filtered by -log10P adj > 1.3, log2 Fold Change > ± 1. Totally genes were 2 genes upregulated whereas 13 genes were downregulated; **(B)** Functional enrichment of GO analysis (Top 10) and signaling pathway enrichment of KEGG analysis (Top 10) were presented to demonstration the potential mechanism of DEGs in comparison of ISO with control group. The red box represents functional/signal pathways associated with ventricular remodeling; **(C)** In comparison of ISO+β-Ecd with ISO group, differential expression genes (DEGs) were analyzed and calculated by Volcano plot. DEGs were then filtered by -log10P adj > 1.3, log2 Fold Change > ± 1. Totally genes were 14 genes upregulated whereas 11 genes were downregulated; **(D)** Functional enrichment of GO analysis (Top 10) and signaling pathway enrichment of KEGG analysis (Top 10) were presented to demonstration the potential mechanism of DEGs in comparison of ISO+β-Ecd with ISO group. The red box represents functional/signal pathways associated with cardiac remodeling and neurotransmitter-related signaling pathways.

To better understand the crosstalk interactions between the central and peripheral systems during sympathetic overactivation induced by ISO, as well as the mechanisms of β-ecdysterone, we conducted weighted correlation network analysis (WGCNA) (Fig. 8A) and delineated the central-peripheral co-molecular mechanisms by which β-ecdysterone improves ventricular remodeling (Fig. 8B). The genes in the "lightcan" module were found to be significantly correlated with cardiac remodeling parameters, including LVHI, EF, FS, fibrotic area, LV mass, HW/BW, inflammatory factor levels of IL-1β, and oxidative stress levels of MDA (Fig. 8B & C). Further exploration involved the gene co-expression network analysis in Cytoscape to identify key genes in the "lightcan" module among multi-organ regulation in the process of cardiac regulation. The gene network (Fig. 8D) highlighted *Dhx37*, an RNA helicase, as a key regulator among the "lightcan" genes. To delve into the mechanism of action of *Dhx37* in cardiac remodeling and its relationship with β-ecdysterone, a literature search was conducted using resources like the Open Map of Knowledge. Up to October 22, 2023, there were 36 relevant studies associated with gender development, cancer, Alzheimer’s disease, RNA modification, and other topics, but none related to cardiovascular research (Fig. S9). As a next step, we analyzed the expression of the *Dhx37* gene in heart and medullary tissue, respectively, and found a significant reduction in *Dhx37* when β-ecdysterone was administered to heart tissue (Fig. S10A & B). In summary, our findings collectively identified the potential role of *Dhx37* in the cardiac central-peripheral crosstalk mechanism, both in the context of ISO-induced cardiac remodeling and the process of β-ecdysterone-treated improvement in cardiac remodeling.

**Figure 8.**
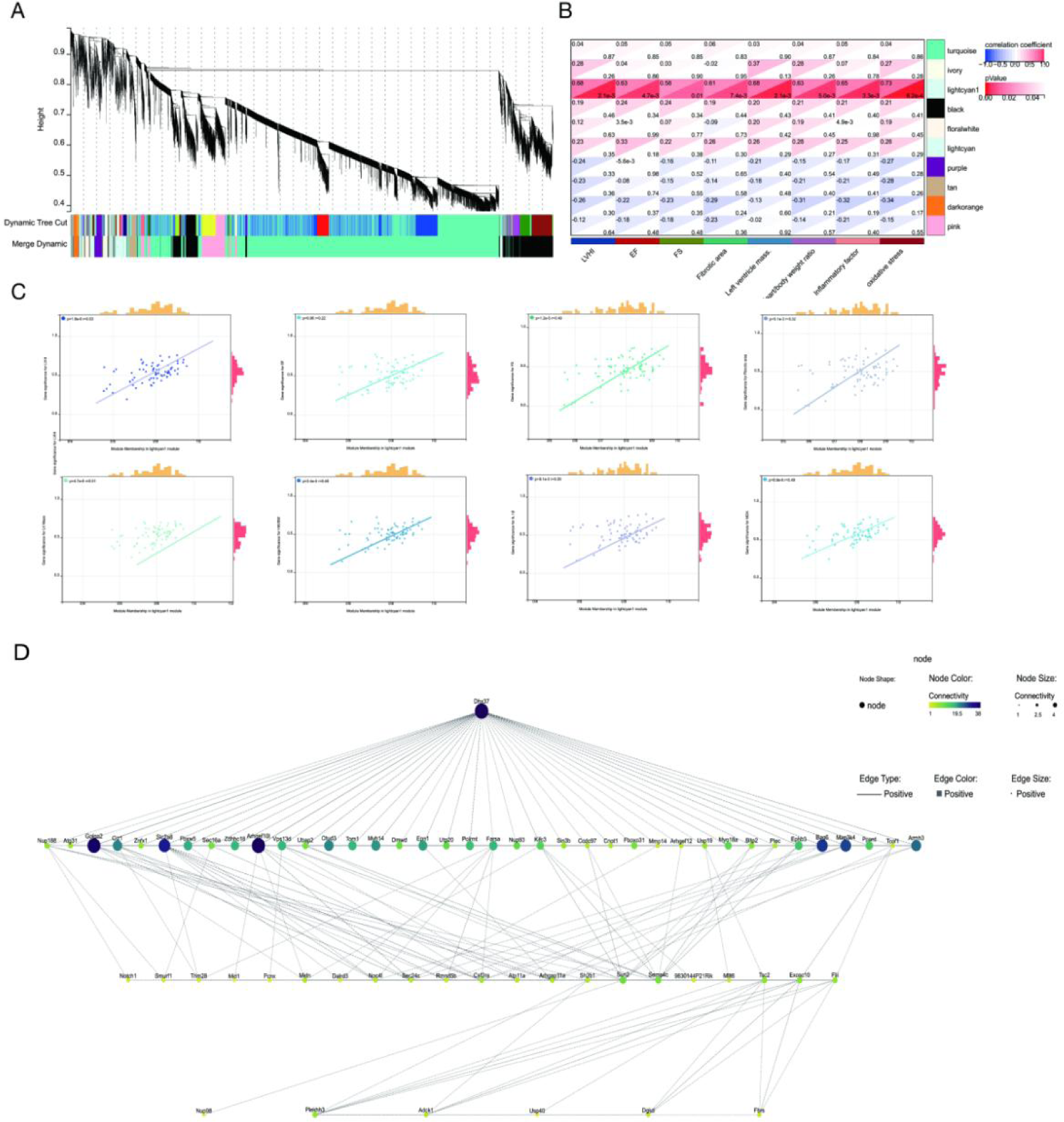
Data Mining for the Correlation between Multi-Organ Transcriptomics of the Heart and Medulla with Ventricular Remodeling Characteristics. **(A)** WGCNA analyses were performed among the transcriptomics of heart and medulla tissue among control, ISO, and ISO+β-ecdysterone groups. n=18; **(B)** Pearson correlation analysis were performed among different module in WGCNA analysis and cardiac remodeling parameters (LVHI, EF, FS, fibrotic area, LV mass, HW/BW, IL-1β and MDA); **(C)** Linear regression correlation analysis were performed with the module membership in lightcyan module and gene significance with cardiac remodeling parameters of LVHI, EF, FS, fibrotic area, LV mass, HW/BW, IL-1β and MDA; **(D)** Network of genes in lightcan module were analysis by Cytoscape. The diagram shows the regulatory relationship among the genes, and *Dhx37* is the most important key regulator presented in this network.

In the final section, to further mining the potential relationship among the aforementioned molecules, including NRF2, of which computer-aided drug design explored β-ecdysterone as a potential competitor, *Hmox-1*, which is considered as the direct downstream factor of NRF2 participating in anti-oxidative stress (Fig. S1C), and Slc41a3, from heart transcriptome data and immunohistochemistry (Fig. S5), a correlation analysis were performed and uncovered a strong and statistically significant positive correlation between *Slc41a3* and *Dhx37* (Fig. S11). By utilizing the UCSC browser and PROMO, it was found that NRF2 is the potential transcription factor for *Slc41a3* and *Dhx37*, respectively (Fig. S12). However, direct evidence clarifying the specific regulatory relationship of NRF2 with *Slc41a3* and *Dhx37* in the process of cardiac remodeling and β-ecdysterone treatment was not available, emphasizing the need for further in-depth mechanistic research based on our findings.

## Discussion

In our study, we made three significant findings: 1) Through computer-aided drug design (CADD), we discovered that the natural MFH bioactive compound β-ecdysterone improved cardiac remodeling by upregulating the NRF2-mediated oxidative stress response; 2) By employing transcriptomics of cardiac and medullary tissue and rs-fMRI techniques in PVN, we found that β-ecdysterone influenced the functionality of the cardiac central nervous system; 3) Through in-depth mechanism mining analysis, we identified the potential cross-talk mechanism involving NRF2, *Slc41a3*, and *Dxh37* in cardiac remodeling and the effects of β-ecdysterone administration in central-peripheral cardiovascular system.

### Medicine and food homology products possess critical therapeutical roles

Throughout history, natural product compounds have played crucial roles in drug research and development. Paclitaxel, extracted from the Pacific yew tree, is a natural product widely employed in treating various cancers, including ovarian cancer, breast cancer^23, 24^, lung cancer, melanoma, lymphoma, and brain tumors^25^. Penicillin, a natural product derived from fungi, finds extensive use in combating infectious diseases such as streptococcal infections^26^, including pharyngitis^27^, tonsillitis^28^, scarlet fever^29^, and more. Data indicates that from 1981 to 2019, 60% of approved anticancer drugs^30–32^ and 75% of approved antimicrobial drugs can be traced back to their natural origins^33^.

Among natural product compounds, medicine and food homology (MFH) products have garnered significant attention, served dual roles as essential dietary components, and possessed pharmacological effects that offer the potential to combat diseases and provide health benefits^34^. For instance, S. grosvenorii extract prevents the generation of superoxide anion and inhibits histamine release from mast cells, thereby reducing histamine-induced nose scratching behavior in ICR mice^35^; Ginsenosides have been found to function as an anti-aging agent by promoting metabolism, protecting the skin and nerves, modulating intestinal flora, and maintaining mitochondrial function^36^; Nardosinone and aurantio-obtusin are recognized as anti-influenza agents^37^; Curcumin extracted from turmeric has been identified for its capacity to alleviate radiation-induced cellular damage, providing a protective shield against radiation harm^15, 38^; Soy isoflavones present in kudzu root exhibit potent free radical scavenging capabilities, mitigating oxidative damage to cells^39, 40^; Lycium barbarum rich in various antioxidants like polysaccharides and anthocyanins, it demonstrates robust cell protection against oxidative stress, endoplasmic reticulum stress, and autophagy-induced apoptosis^41^; Ascorbic acid (Vitamin C) serves as a formidable antioxidant, neutralizing free radicals, reducing oxidative damage from UV-Vis irradiation^42^, corneal endothelial damage^43^, and fostering efficient cell repair and regeneration, etc.

### Computer-based approaches facilitate natural drug discovery with high efficiency and accuracy

The traditional exploration of potential pharmacological effects of natural product compounds, as seen in the investigation of cardiovascular diseases, often adopts a research focus centered on a concept. While this approach proves valuable in uncovering the multifaceted nature of natural products, extending their pharmacological effects, e.g. from cancer to cardiovascular diseases, lacking a sufficiently robust research foundation. To address these research limitations, we currently have implemented a series of comprehensive approaches to establish a solid research foundation.

First, through bioinformatic and literature research, we initially focused on the key signaling pathway in cardiac remodeling: oxidative stress, and its major regulatory factor NRF2. The activation of NRF2 is crucial for maintaining the oxidative/antioxidative balance^44, 45^. Activation of NRF2 has been described in numerous pathologies, including cancer, cardiovascular, respiratory, renal, digestive, metabolic, autoimmune, and neurodegenerative diseases^46^. For instance, dysregulated NRF2 confers high-level resistance to anticancer drugs and reactive oxygen species (ROS) on cancer cells and directs cancer cells toward metabolic reprogramming^47^. Considering the increasing interest in discovering novel NRF2 activators due to its clinical application, our efforts were devoted to the development of MFH compound able to induce NRF2 nuclear accumulation by targeting its natural repressor protein Kelch-like ECH-associated protein 1 (KEAP1) through covalent modifications on cysteine residues^48^, the combination of which inhibits the transfer of NRF2 to the nucleus through ubiquitination and subsequent degradation of NRF2.

Second, we used the KEAP1-NRF2 binding site as the starting point, designed the competitive binding pharmacophore through utilizing CADD approaches among medicine and food homology compound database (MFCD), which was first concluded and integrated with its available composition and structure by our study, containing 3907 compounds from 110 MFH products. CADD is an emerging field that has drawn a lot of interest because of its potential to expedite and lower the cost of the drug development process. Drug discovery research is expensive and time-consuming, often taking 10-15 years for a drug to be commercially available. CADD has significantly impacted this area of research^49^. CADD techniques have garnered a great deal of attention in academia and industry due to their versatility, low costs, possibilities of cost reduction in *in vitro* screening, and in the development of synthetic steps; these techniques are compared with high-throughput screening, particularly for candidate drugs. Thus, CADD approaches play a significant role in the drug development process since they allow better administration of resources with successful results and a promising future market and clinical-wise^50^. The most significant achievement of CADD is the discovery of Paxlovid, the first orally available COVID-19 drug^51^, which contributes to the structure-based drug design, providing the most available approach in facing present and future pandemics. Through CADD approaches, researchers have provided useful evidence to discover anti-inflammatory drugs towards TNF-α, which is associated with immune diseases, cancer, and psychiatric disorders^19^; Similarly, by using CADD approaches, anti-Alzheimer drugs^52^, anti-diabetic therapies^53^, and anti-cancer agents^54, 55^ have been rapidly screened and evaluated. Based on CADD approaches, which aims to bridge the research deficiencies by combining experimental and computational methodologies, providing a more systematic and informed exploration of the pharmacological potential of natural products, specifically in the context of cardiac remodeling and oxidative stress regulation, we selected the best promising compound in MFCD, β-ecdysterone, with high potential to combine with KEAP1-NRF2 and qualified drug-likeness.

### β-ecdysterone, the promising functional compound with multiple pharmacological effects

Notably, the studies on β-ecdysterone have predominantly focused on bone-related research. Researchers have revealed that β-ecdysterone alleviates osteoarthritis by activating autophagy in chondrocytes through the regulation of the PI3K/AKT/mTOR signaling pathway^56^. Another study investigated the regulation of osteoblast autophagy based on the PI3K/AKT/mTOR signaling pathway, demonstrating the effect of β-ecdysterone on fracture healing^57^. Additionally, β-ecdysterone has been shown to enhance bone regeneration through the BMP-2/SMAD/RUNX2/Osterix signaling pathway^58^. Moreover, the beneficial effects of β-ecdysterone have been gradually addressed, including its therapeutic potential in hemodynamics and myocardial contractility on coronary artery occlusion-induced arrhythmias^59^. Therefore, based on previous studies and our current research, β-ecdysterone emerges as a beneficial active ingredient with therapeutic value. Through our isoproterenol (ISO)-induced animal model and AngII-induced cell model, β-ecdysterone exhibited a series of effects on ameliorating cardiac remodeling and oxidative stress. Interestingly, our findings revealed that the anti-hypertrophic effects on cardiomyocytes were more pronounced than the anti-fibrotic effects on cardiac fibroblasts when treated with β-ecdysterone. This observation was evident in terms of fold changes, concentration-dependent responses, effective dose, as well as NRF2 expression and oxidative stress. In the cytoplasm, NRF2 binds with KEAP-1, undergoes ubiquitination, and subsequently degrades, leaving a portion of NRF2 waiting in the cell nucleus. Under AngII stimulation, cardiac myofibroblasts are activated, accelerating cell proliferation, and increasing oxidative stress levels. Consequently, the content of NRF2 in the cytoplasm of cardiac myofibroblasts increases, which is consistent with our Figure 3C. According to the result of the oxidative stress metabolite MDA in cardiac fibroblast, it shows a less excessive accumulation of oxidative stress substances in cardiac myofibroblasts, indicating the balance achieved at the appropriate oxidative stress levels. These findings may explain that the beneficial effects of β-ecdysterone on NRF2 may not be as pronounced in cardiac myofibroblasts as they are in cardiomyocytes.

### Interdisplinary investigation of central-peripheral cardiovascular crosstalk regulation contribute to in-depth insight of etiological mechanisms and therapeutical targets

Cardiovascular system involves a feedback mechanism between the central and peripheral nervous systems. There remained a question of whether the administration of β-ecdysterone affects the activity of the central sympathetic nervous system. According to the results of the cardiac transcriptome, it is intriguing that in the treatment of β-ecdysterone, many alterations in neurotransmitter-related signaling pathways were observed in heart tissues. Based on the drug-likeness evaluation in Table S3, β-ecdysterone is unable to pass through the blood-brain barrier and cannot directly act on the central nervous system. Since our pathological cardiac remodeling model was established via the method of sympathetic overactivation by ISO^21, 60, 61^, overactivated sympathetic nerves cause dysregulation of the central nervous system and disrupt the balance of activities between sympathetic and parasympathetic nerves^62^. To further explain why the treatment of β-ecdysterone caused alterations in neurotransmitter-related signaling pathways, we designed the following interdisplinary approaches to investigate the crosstalk mechanism of central-peripheral regulation.

First, we measured neurotransmitters, including norepinephrine (NE) and acetylcholine (Ach), in heart tissue, along with 33 other neurotransmitters, and revealed that β-ecdysterone significantly decreased NE and its metabolites, indicating its inhibitory effect on sympathetic activity. This suggests that β-ecdysterone counteracts sympathetic overactivation without notable impact on parasympathetic neurotransmitters in ISO-induced cardiac remodeling.

Second, we explored whether β-ecdysterone treatment affects the sympathetic central nervous system. We applied resting-state functional magnetic resonance imaging (rs-fMRI) to achieve real-time and non-invasive detection of whole brain activity^62–64^. Animal studies using rs-fMRI are often applied to nervous system diseases. For instance, regional homogeneity alterations of rs-fMRI were used to examine chronic rhinosinusitis with olfactory dysfunction^65^, mapping the aberrant brain functional connectivity in new daily persistent headache^66^, or applied in revealing the aberrant large-scale network interactions across psychiatric disorders^67^. Given these rs-fMRI experiences, in our current study, we found that the sympathetic nerve center PVN showed inhibitory activation when treated with β-ecdysterone compared with the ISO group, ensuring the effects of β-ecdysterone on the central nervous system and expanding the pharmacological effects of β-ecdysterone beyond bone regeneration^56, 58^, osteoporosis^65^, osteoarthritis^56^, and kidney injury^66^. In addition to the sympathetic nerve center PVN, the medulla oblongata is the center of the cardiovascular system^68^, containing cardiac vagal neurons and sympathetic neurons.

Third, to further explore the possible central mechanism and its association with the characterization of cardiac remodeling, we applied a series of data analysis methods, including WGCNA analysis of peripheral cardiac tissue and medullary tissue transcriptomic data with correlationship of cardiac remodeling characterization. The significance of transcriptome research lies in gaining a deeper understanding of the effects and regulatory mechanisms of drugs at the gene level. It facilitates the unraveling of molecular mechanisms of drug action, prediction of drug responses and individual differences, and consequently, the anticipation of drug efficacy and variations in individual responses. Additionally, it aids in the discovery of new drug targets and biomarkers. Our study discovered cardiac central-peripheral regulation in the β-ecdysterone-treated ISO group, seeking related genes and their key regulator, *Dhx37*, through data processing. *Dhx37* is an RNA helicase, and existing studies have associated it with liver cancer progression by cooperating with PLRG^69^. Mutations and dysregulation of *Dhx37* impact the prognosis of hepatocellular carcinoma and lung adenocarcinoma^70^. Although research has mainly focused on its potential as a carcinoma biomarker and its mechanistic function as an RNA helicase in the release of the U3 snoRNP from pre-ribosomal particles^71^, and activation by UTP14A in ribosome biogenesis^72^, our study creatively found the potential role of *Dhx37* in cardiovascular disease.

Additionally, through correlation analysis, the potential association of the key genes involved in this study was explored. The results showed that *Dhx37* was highly positively correlated with *Slc41a3*. However, the precise function of *Slc41a3* in the heart remains incompletely understood, as research in this area is still in its early stages. Insights gleaned from the examination of *Slc41a3* in other tissues offer potential implications: 1) Ion Balance: *Slc41a3* participates in the regulation of ion balance in cardiac cells, including ions like calcium, magnesium, or zinc^73^. These ions play crucial roles in the normal functioning of cardiac muscle and the electrophysiology of the heart. 2) Cardiac Contraction: Certain ions, such as calcium, play critical roles in the contraction and relaxation of cardiac muscle. *Slc41a3* may influence cardiac contraction by its regulatory function on magnesium homeostasis^74^ and is positively correlated with magnesium levels in serum^75^, suggesting that it may act as a magnesium extrusion mechanism across either the plasma membrane or organellar membranes^76, 77^. *Slc41a3* was also found to be involved in the regulation of pulmonary hypertension^78^ and diabetes mellitus^79^. Similar to *Dhx37*, *Slc41a3* is also considered carcinoma-related, with upregulation identified as a liver hepatocellular carcinoma biomarker^80^ associated with poor prognosis^81^.

To conclude, our study provides a novel regulatory possibility: NRF2-regulated *Slc41a3*/*Dhx37*, identified through promoter transcriptional prediction. The results showed that NRF2 could bind to both the promoter of *Slc41a3* and *Dhx37*, regulating their expression levels (Figure S12). Unfortunately, no research has investigated the relationship among NRF2, *Slc41a3*, and *Dhx37* in any type of cells or organs, presenting significant investigatory potential to be further illuminated.

### Perspectives and limitations

For the perspectives, as a study in the field of high-throughput, broad-spectrum natural product discovery and mechanism exploration, we aimed to use macro-level screening methods to focus on a specific compound and its potential mechanisms. Our interdisciplinary research merged computer science and pharmacology, pinpointing the natural MFH compound β-ecdysterone as an NRF2 enhancer in cardiac remodeling. Exploring cardiac transcriptome data unearthed β-ecdysterone impact on peripheral neurotransmitters, confirmed through neurotransmitter metabolomics, rs-fMRI, and medullary transcriptome analysis. Delving into crosstalk mechanisms, we identified *Slc41a3* as a key regulator influenced by β-ecdysterone, and WGCNA linked multi-organ transcriptomics to cardiac remodeling, revealing pivotal genes like *Dhx37*. Further investigation into regulatory relationships unveiled a positive correlation between *Slc41a3* and *Dhx37*. Predicting NRF2 as the transcription factor for *Slc41a3* and *Dhx37* suggests novel directions for future in-depth studies on regulatory mechanisms in anti-oxidative stress processes. The whole process of our functional MFH compound and in-depth data mining was depicted as Figure S13.

However, this approach has its limitations. For instance, no mechanism research and verifications were performed among NRF2, *Slc41a3* and *Dhx37*, as our research objective was set for data mining. Despite these limitations, this method holds significant research and scientific value. With the expansion of this research, more researchers can concentrate on such research methods, thereby broadening the scope of early target-compound screening. Therefore, the methodology employed in this study carries significant practical value. We firmly believe that the continuity of research is of utmost importance. It is not only about discovering new phenomena but also about providing clues for future investigations.

In conclusion, our research yields valuable insights of functional compound β-ecdysterone of MFH through high-throughput and efficient computational methods, complemented by validation of its beneficial effects on cardiac remodeling using animal and cell models. It significantly contributes to a deeper understanding of crosstalk mechanism between cardiac central and peripheral systems, providing novel insights into the potential application of natural MFH functional products in the treatment of cardiovascular diseases.

## Acknowledgements

We thank Dr. Yanyan Zhang and Dr. Weiwei Jia for TAC and L-NAME animal models. We thank Dr. Yu Jiao and Dr. Xiaoqing Qin for performing rs-fMRI and data analysis.

